# Structured illumination microscopy artifacts caused by illumination scattering

**DOI:** 10.1101/2021.01.01.425033

**Authors:** Yanquan Mo, Fan Feng, Heng Mao, Junchao Fan, Liangyi Chen

**Affiliations:** State Key Laboratory of Membrane Biology, Beijing Key Laboratory of Cardiometabolic Molecular Medicine, Institute of Molecular Medicine, School of Future Technology, Peking University, Beijing 100871, China; Center for Bioinformatics, National Laboratory of Protein Engineering and Plant Genetic Engineering, School of Life Sciences, Peking University, Beijing 100871, China; School of Mathematical Sciences, Peking University, Beijing 100871, China; Chongqing Key Laboratory of Image Cognition, College of Computer Science and Technology, Chongqing University of Posts and Telecommunications, Chongqing 400065, China; PKU-IDG/McGovern Institute for Brain Research, Beijing 100871, China; Beijing Academy of Artificial Intelligence, Beijing 100871, China; Shenzhen Bay Laboratory, Shenzhen 518055, China

**Keywords:** structured illumination microscopy, aberration, illumination path, optical scattering

## Abstract

Despite its wide application in live-cell super-resolution (SR) imaging, structured illumination microscopy (SIM) suffers from aberrations caused by various sources. Although artifacts generated from inaccurate reconstruction parameter estimation and noise amplification can be minimized, aberrations due to the scattering of excitation light on samples have rarely been investigated. In this paper, by simulating multiple subcellular structure with the distinct refractive index (RI) from water, we study how different thicknesses of this subcellular structure scatter incident light on its optical path of SIM excitation. Because aberrant interference light aggravates with the increase in sample thickness, the reconstruction of the 2D-SIM SR image degraded with the change of focus along the axial axis. Therefore, this work may guide the future development of algorithms to suppress SIM artifacts caused by scattering in thick samples.

## Introduction

By selectively highlighting structures with fluorescent proteins or dyes, fluorescence microscopy achieves high contrast and resolution and has been widely used in biomedical research. Super-resolution (SR) fluorescence microscopy techniques, including stimulated emission depletion (STED)[1], photoactivated localization microscopy (PALM)[2,3], stochastic optical reconstruction microscopy (STORM)[4], and structured illumination microscopy (SIM)[5,6] emerged at the beginning of the 21st century. Using different principles, they have overcome the physical diffraction limit and enabled visualization of many previously unappreciated, intricate cellular structures and organelles. Among these techniques, SIM excels in extending the spatial resolution with a relatively small increase in emission photons [7]. Thus, SIM is suitable for live-cell SR imaging, such as identification of vesicle fusion intermediates including enlarged fusion pores by 300 Hz SR imaging, one-hour time-lapse SR imaging of actin filaments labeled by lifeact-EGFP with minimal photobleaching[8], and 2,000-frame continuous mitochondrial SR imaging with small enlargements of cristae diameters[9].

Despite these advantages, SIM is prone to reconstruction artifacts[10]. Artifacts due to incorrect reconstructed parameters can be suppressed by designing algorithms to determine parameters of post image acquisition more accurately [8,11–14]. Meanwhile, reduced photon dosage yields low signal-to-noise contrast in the raw images captured, which manifests artifacts as the amplification of noise by the Wiener inverse filtering process. By introducing the *a priori* continuity of any structures along space and time as the penalty term based on the Hessian matrix, these intrinsic artifacts can be effectively reduced[8]. In addition to artifacts generated due to the reconstruction algorithms, an imperfection in the optical path causes aberrations that significantly distort the reconstructed SR images[15]. For example, Débarre et al. classified aberrations of the light path of structured illumination into two groups: those that affected the illumination pattern of excitation and those that did not [16]. Thomas et al. demonstrated SIM through 35 *μ*m of *Caenorhabditis elegans* tissue using adaptive optics (AO) to correct aberrations, which resulted in images with a resolution of 140 *nm* [17]. Žurauskas et al. presented IsoSense, a wavefront sensing method that mitigated sample dependency in image-based sensorless adaptive optics applications in microscopy. They demonstrated the feasibility of IsoSense for aberration correction in a deformable-mirror-based SIM [18]. By combining AO with SIM imaging, Lin et al. presented an improved resolution (140 *nm* laterally and 585 *nm* axially) and reduced artifact imaging in 3D SR imaging [19]. Liu et al. simulated the effects of the biased thickness of coverslip, the tilted coverslip, the mismatched refractive index (RI), and the misalignment of incident beams concerning the back focal plane of the objective[20].

Most of these previous attempts focused on correcting aberration on the emission detection path of SIM systems. Some work has also analyzed and corrected the aberrations of light paths within the microscope, such as IsoSense. On the other hand, more detailed analysis regarding the SIM illumination path, such as the effect of scattering and absorption of the complex thick sample on the illumination pattern are still required. SIM depends on tightly focused lights that interfere with generating the grid-patterned illumination, which would be destroyed in a deep sample. This issue is best demonstrated in **Fig. 1A**: when the incident light illuminates the sample, scattering occurs and produces aberrations due to the inhomogeneous sample with a different RI. Due to the aberrations, the illumination pattern projected inside the sample becomes distorted, which will distort the reconstruction of SR images to misleading results.

**Fig. 1.**
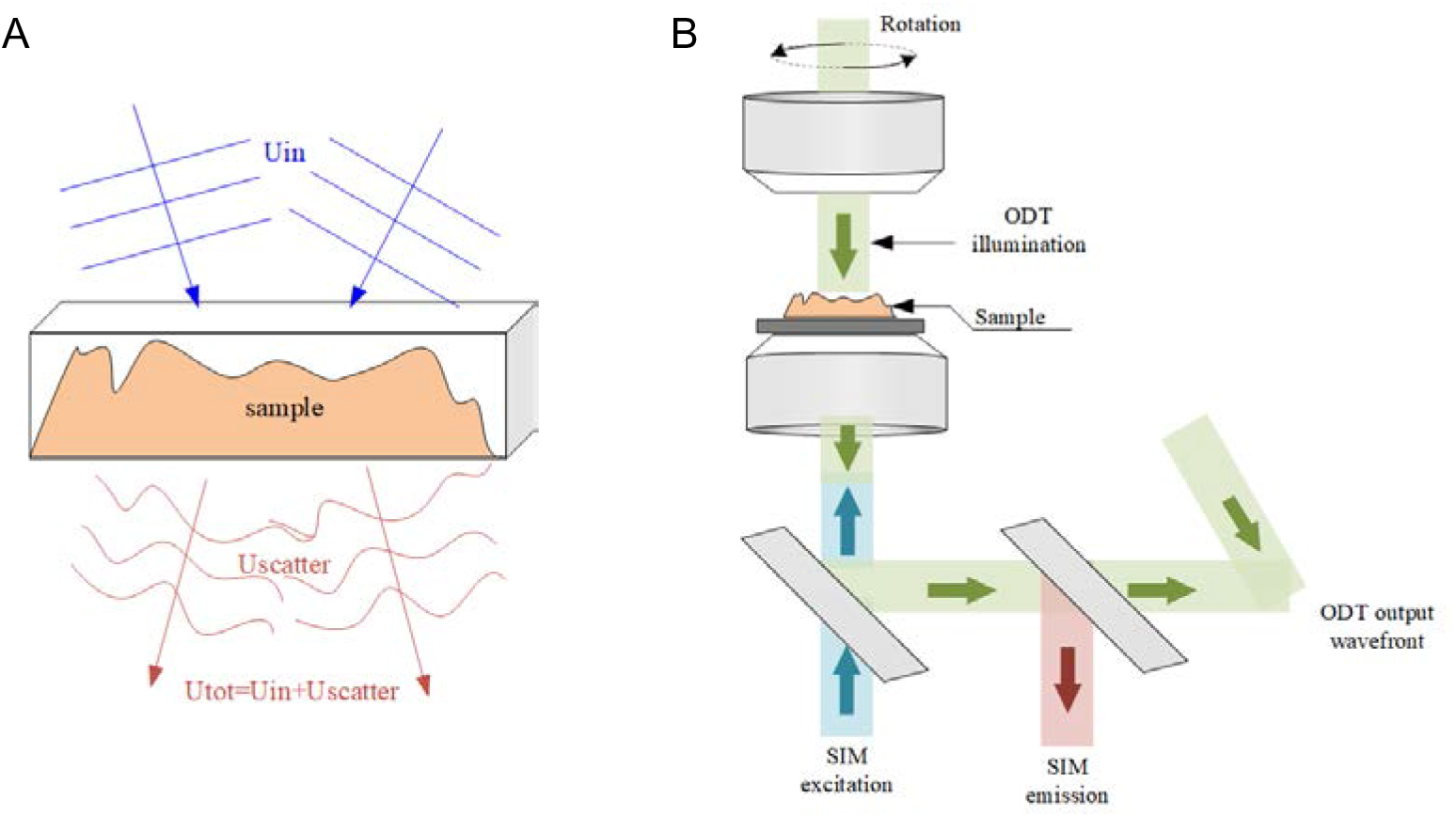
Schematic diagram of incident light scattering in a thick sample and the SR-FACT optical system. (A) Process in which incident light scatters and produces aberrations in an inhomogeneous thick sample. (B) Dual-mode SR-FACT optical system which can acquire the three-dimensional refractive index distribution of the entire thick sample.

Dong et al. developed SR fluorescence-assisted diffraction computational tomography (SR-FACT), which combines label-free three-dimensional optical diffraction tomography (ODT) with two-dimensional fluorescence Hessian SIM [21]. The optical path diagram is shown in **Fig. 1B**. The dual-mode uses the same detection objective lens to image the same cell at the same location. ODT can acquire the three-dimensional RI distributions of the entire cell in these two modes, which reveal organelles with different refractive indices, such as lipid droplets, mitochondria, and lysosomes[21]. Therefore, even in samples such as live cells, it is expected that organelles with different RI may scatter and distort the incident light, which causes reconstruction artifacts. Meanwhile, since light scattering is closely related to the RI distribution of the sample, we can calculate the scattering and distortion of the SIM incident light using the RI yielded by the ODT.

In this paper, we built a model with preset RI distribution of the structure and calculated SIM incident light scattering. Consequently, we could obtain distortions of the illumination in sinusoidal patterns and analyzed the effect which these distortions have on the reconstructed SR images.

## Method

In SIM, the emission distribution *D*(***r***) detected by the camera can be expressed as:

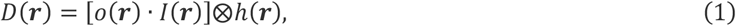

where *o*(***r***) is the spatial distribution of the object labeled with fluorophores, *I*(***r***) is the sinusoidal intensity pattern for illumination, *h*(***r***) is the point spread function (PSF) of the detection path, and ⊗ is the convolution operator. ***r*** ≡ (x, y, z) is the space position vector; in general, z is ignored due to imaging at the focal plane in 2D SIM. The illumination of 2D SIM is commonly composed of 9 sinusoidal patterns in 3 orientations and 3 phases; therefore, *I*(***r***) can be described by the following equation:

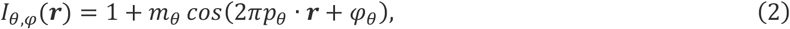

where *m*_*θ*_ is the modulation depth, ***p***_*θ*_ is the pattern period, and *φ*_*θ*_ is the initial phase in orientation *θ*. In the Fourier domain, equation (2) is expressed as:

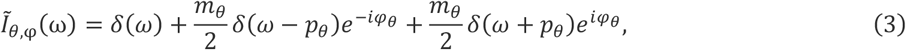

where 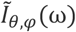 is the Fourier transform of *I*_*θ,φ*_(***r***), *δ*(*ω*) is the Dirac function, and ω is the spatial frequency. According to equations (1) and (3), we obtain the Fourier spectrum of the detected image:

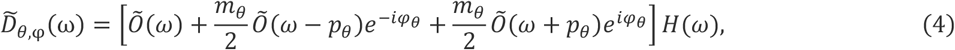

where 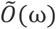 is the spectrum of an object, and 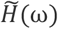 is the optical transfer function (OTF) of the optical system. Equation (4) demonstrates how the high spatial frequency information is shifted into the observable region of the microscope OTF. 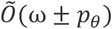 is the ±*p*_*θ*_ frequency shift of 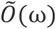 and contains high-frequency components that are normally outside of the OTF. Finally, after combining the extracted high-frequency information with equation (4) under 9 illumination patterns of different orientations and phases, the SR SIM image can be obtained by deconvolution.

Based on SIM’s forward model and reconstruction process, we can analyze the incident light’s scattering process and distortion of illumination patterns in the thick sample. First, we calculate the light scattering field of the incident light. According to Maxwell’s electromagnetic theory, the light propagated at the space coordinate ***r*** satisfies the following wave equation at time *t*:

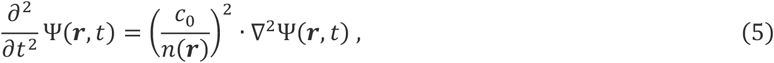

where 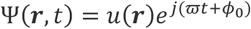 is the spatial and temporal distribution of wave fields, *c*_0_ is the speed of light in the vacuum, *n*(***r***) is the refractive index spatial distribution, ∇^2^ is the Laplace operator, and *u*(***r***) is the plane wave. Next, the Helmholtz equation can be expressed as:

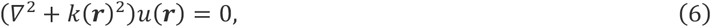

where *k*(***r***) is the distribution of the spatial wave vector. For an inhomogeneous thick sample in the volume, we assume that the averaged wave vector of light in the medium surrounding the sample is *n*_*m*_, the wavenumber is 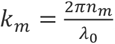, and *λ*_0_ is the wavelength of the incident laser. Therefore, equation (6) can be rewritten as:

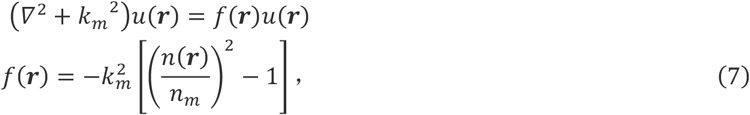

*f*(***r***) is also called the scattering potential, reflecting the heterogeneous distribution of the refractive index in the sample. Using the Green function to solve equation (7), we derive the Lippmann-Schwinger integral formula:

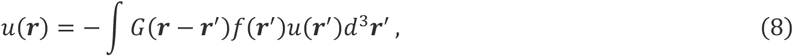

where *G*(***r***) is the Green function:

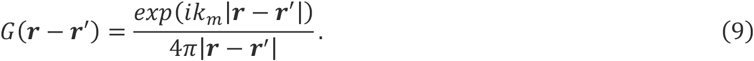

Total output wavefield *u*(***r***) is considered the sum of the incident wavefield *u*_*in*_ (***r***) and scattered wavefield *u*_*s*_ (***r***):

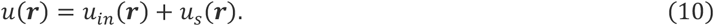

Finally, by solving equation (8), the complex amplitude *u*_*s*_(***r***) generated by the sample scattering, and the subsequent total light field *u*(***r***) can be obtained. We used the Lippmann-Schwinger model (LS model)[22–24] to calculate the scattered field *u*_*s*_(***r***) for the incident illumination in different orientations and phases. The LS model is a superior nonlinear forward model to approximate models such as Born or Rytov, while it can generate accurate estimations. The LS model formula is as:

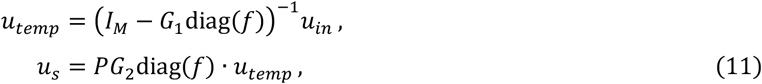

where *I*_*M*_ is the identity matrix, *G*_1_ is the discrete counterpart of the continuous convolution with the Green function for the thick sample volume, and *G*_2_ is the discrete Green function at the measurement plane, diag(*f*) represents the diagonal matrix formed out of the entries of *f*. *P* models the effect of the pupil function of the microscope, and can also encode the contribution of a free-space propagation of light to account for an optical refocus of the measurements.

In the illumination path of SIM, the excitation pattern was formed by the interference of two beams. The complex amplitude distribution of total incident light *u*_*in*_(***r***) was obtained by adding two beams as follows:

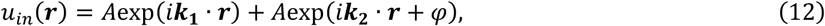

where *A* is the amplitude of two coherent beams, ***k***_1_ and ***k***_2_ are the wavevectors, and *φ* is the phase difference between the two beams that produce illumination patterns with different phases. Therefore, we show the flow chart of the whole simulation process in **Fig. 2**.

## Results

As shown in **Fig. 3A**, we simulated a cuboid with 25.6 *μm*×25.6 *μm*×7 *μm* in size, which was filled with the medium with a RI of *n_m_*=1.33. Then we placed a 3D cell phantom inside the cuboid with different structures with RI between 1.33 and 1.5 (**Fig. 3B**), which is designed to simulate organelles such as lipid droplets or nucleus[25]. We assumed that the focal plane of the SIM imaging is placed at the bottom of the cube, i.e., the incident light must pass through the medium and cell phantom before illuminating the fluorescently labeled sample at the focal plane. The whole sample was composed of the cuboid, cell phantom and ground truth (**Fig. 3C)**. By adjusting thickness *h* of the cell to 0, 2, 3, 4, 5, 6, and 7 *μm*, we simulated the change in the distortion of the illumination by the scatter of different thicknesses. We set the incident wavelength to 532 *nm.* The intensity of *u*_*in*_ (*x*, *y*, *z* = 7*μm*) is the pattern of undistorted SIM illumination at the focal plane with three orientations of the excitation patterns in **Fig. 3D**, where the maximum and minimum intensities of sinusoidal patterns are 1 and 0, respectively. The scattering potential *f*(***r***) can be calculated according to equation (7) based on the size and RI of the simulated cuboid and cell. The intensities of scattered fields *u*_*s*_(*x*, *y*, *z* = 7*μm*) of a single phase in the vertical orientation of the illumination pattern in different *h* are shown in **Fig. 4A.** According to the calculated results, the main direction of the scattered field distribution is along the direction of the incident light. In other words, the incident light at a tilt of 0, 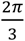, or 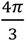 causes the predominant scattering with the same tilt at 0, 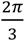, or 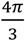. **Fig. 4B** shows the maximum intensity of *u*_*s*_ (***r***) with different *h* in **Fig. 4A**, which demonstrates a nonlinear increase in scattered field intensity when *h* increases. **Fig. 4C** and **4D** are the amplitudes and phases of *u*_*s*_(***r***) at different image planes, which demonstrates intricate distributions of scattered field laterally and axially. The reason is for a thicker sample, the optical path among different incident lights more substantially diverges.

**Fig. 2.**
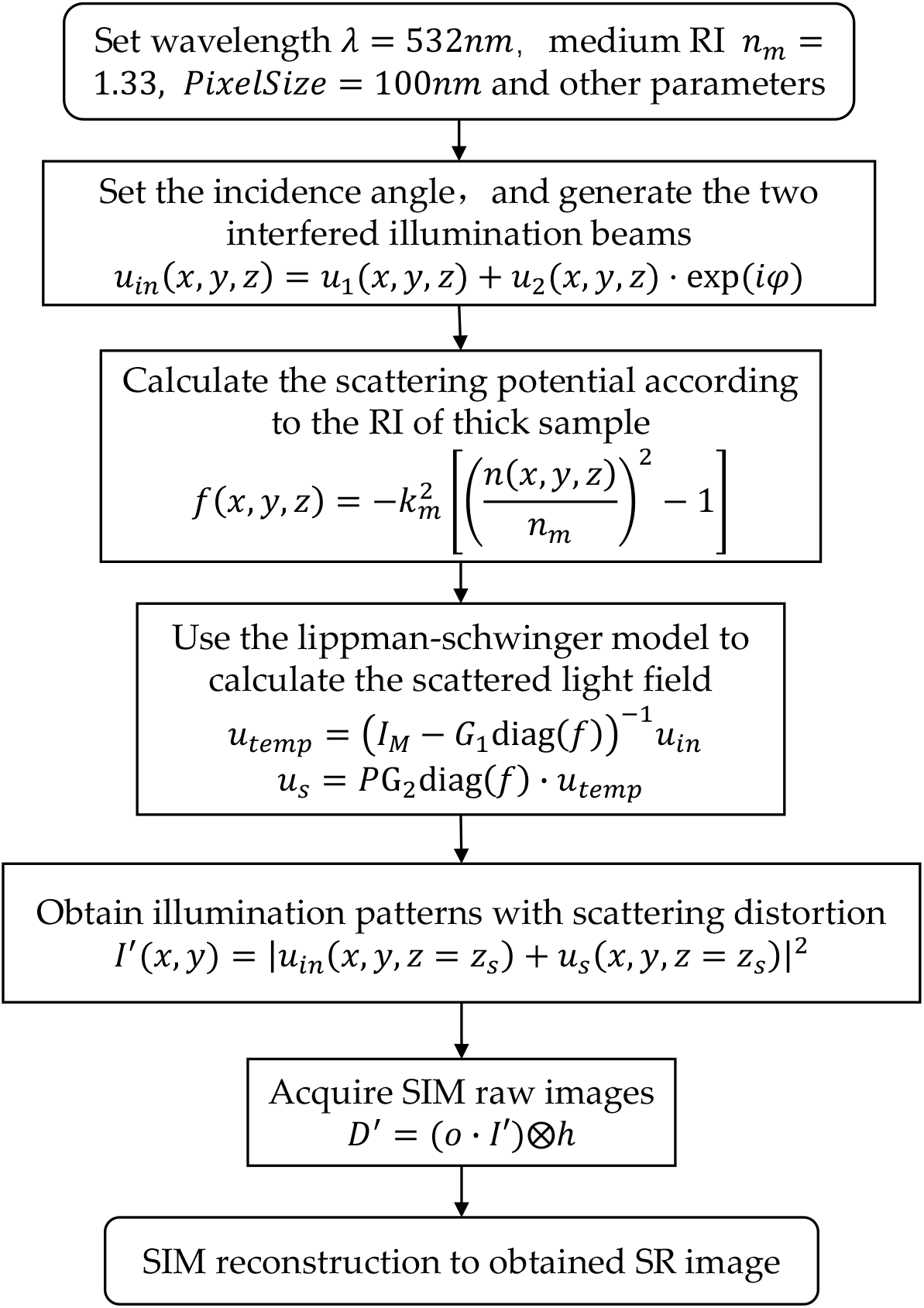
Flow chat of the whole simulation process.

**Fig. 3.**
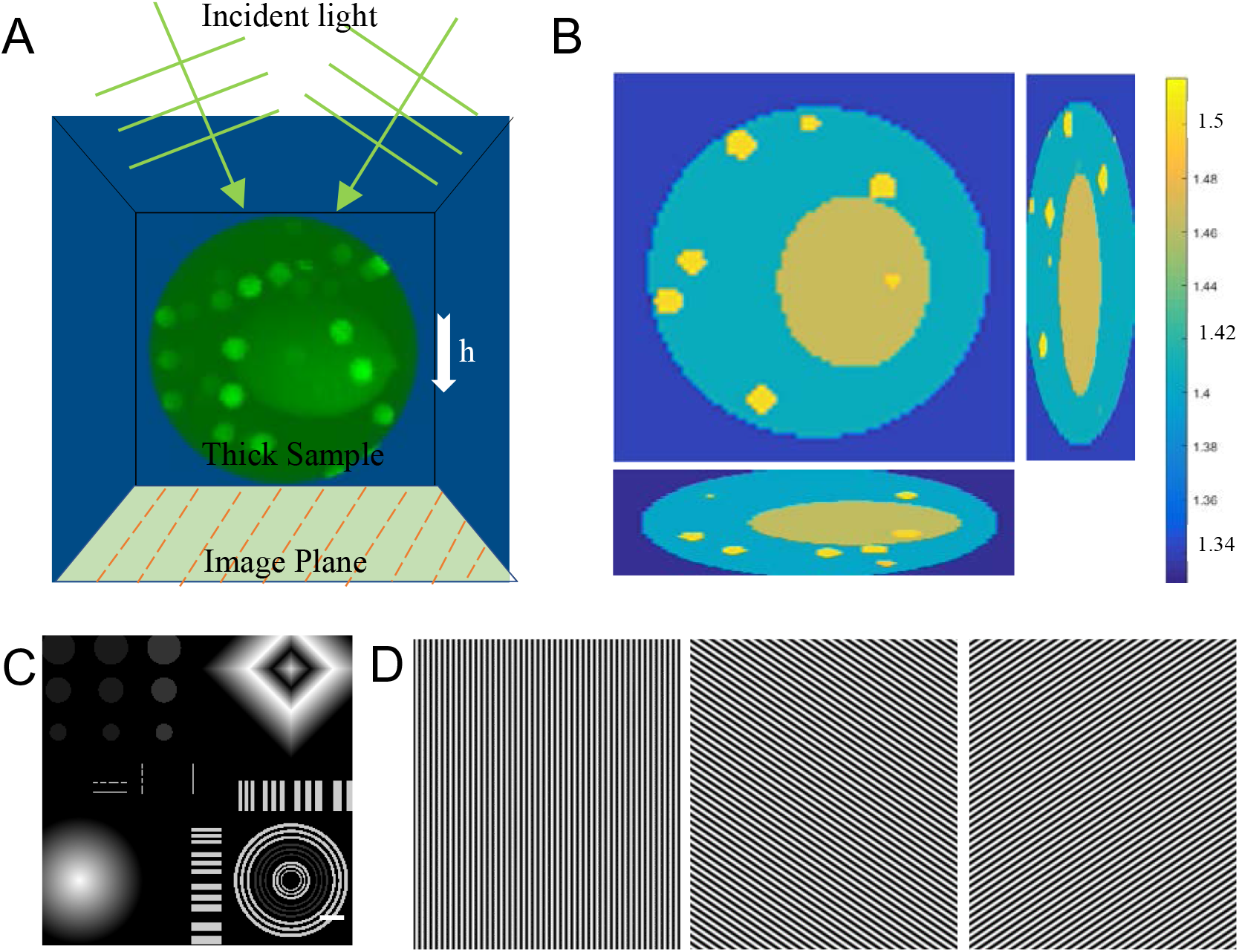
Design of the model and ground-truth sample. (A) Diagram of the forward model of SIM imaging including light scattering in illumination path and imaging. Set the thick sample’s height *h* = 0, 2, 3, 4, 5, 6, and 7 *μm* for 7 different variable values in total. (B) The thick sample’s RI distribution at 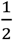 height, 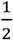 length, and 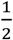 width, respectively. And the RI varies from 1.33 to 1.5, the volume is 25.6 *μm*×25.6 *μm*×7 *μm*. (C) The ground truth of SIM imaging to simulate fluorescence emission distribution. The ground-truth image is the focal plane at the lowest level of the thick sample. Scale bar: 2 *μm*. (D) Sinusoidal excitation patterns in 3 orientations with intensities of 0-1.

**Fig. 4.**
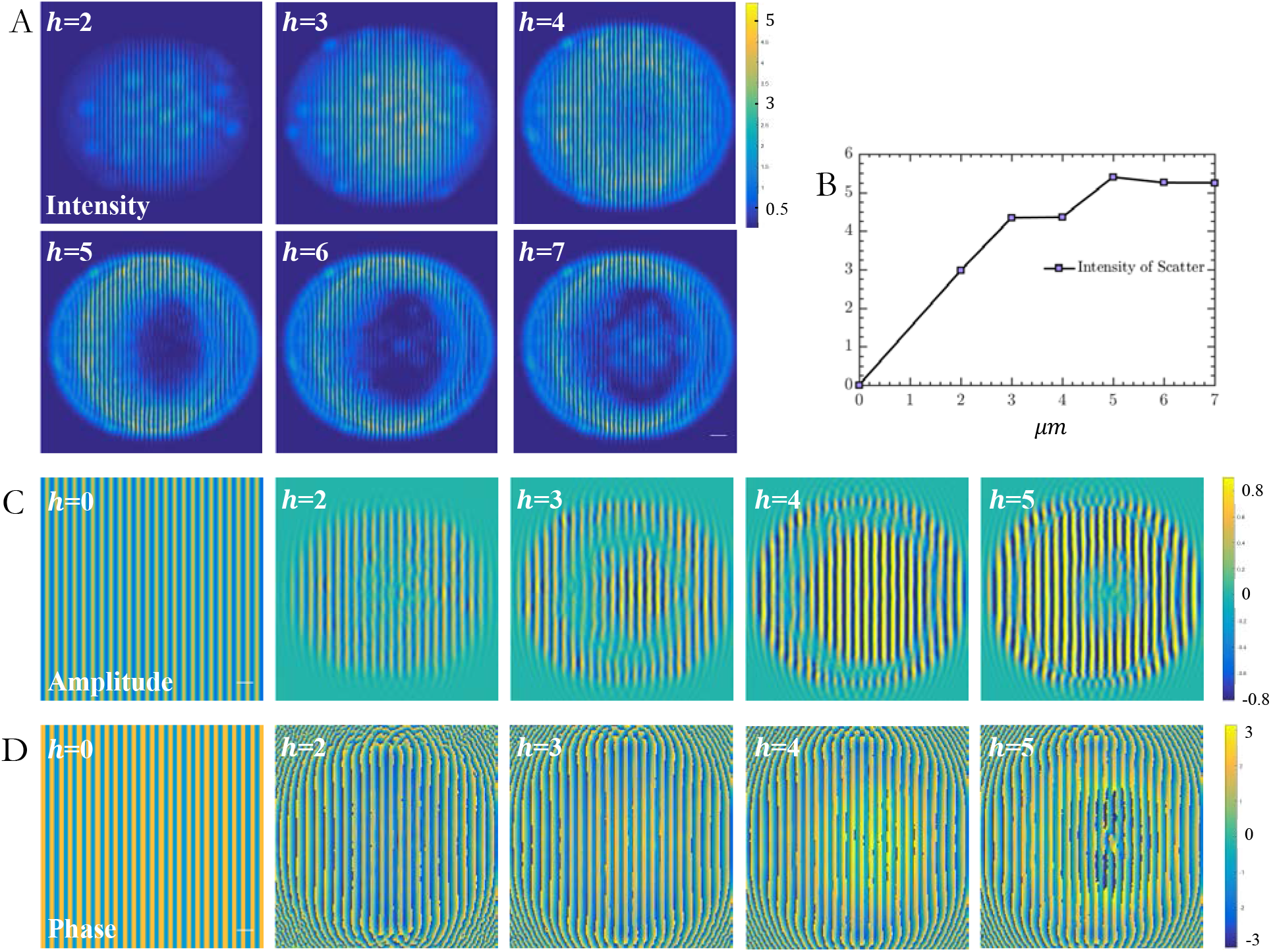
Intensities, amplitudes, and phases of scattered fields in different sample thicknesses. (A) Intensities of the scattered field *u*_*s*_(***r***) of a single phase in the vertical orientation of illumination pattern in different sample thicknesses. (B) Maximum intensity of *u*_*s*_(***r***) in different *h* in (A). (C) Amplitudes of *u*_*s*_(***r***) in different *h*. (D) Phases of *uss*(***r***) in different *h*, which is used to make a comparison with incident light. Scale bar: 2 *μm*.

Having obtained the scattered field’s spatial distribution, we can calculate the distortions of illumination patterns in the focal plane. As shown in **Fig. 5A**, distortions of illumination became severe upon an increase of the image depth. By calculating the structural similarity (SSIM) index of distorted patterns with the non-distorted illumination pattern, we quantitatively analyzed scattering effects axially. The SSIM of the reconstruction results appeared to be negatively correlated with the plane thickness (**Fig. 5B**), which indicated an increased distortion along the axial axis. The SSIM was calculated as follow:

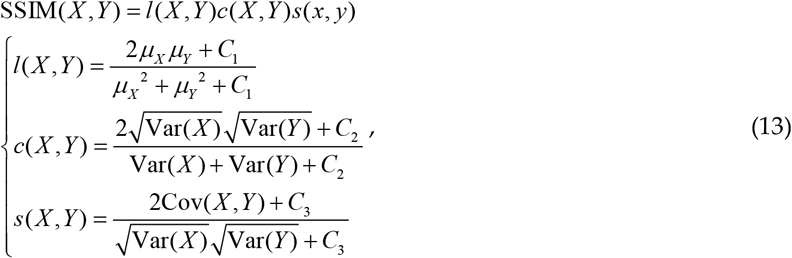

where X, Y represent different images, and Var and Cov are the variance and covariance of all pixels in the designated image respectively. *l*(*X*, *Y*) is the intensity comparison function to measure the similarity of the mean intensities of two images (*μ*_*X*_, *μ*_*Y*_); *c*(*X*, *Y*) is the contrast comparison function to measure the similarity of the contrast of the two images; *s*(*X*, *Y*) is the structure comparison function to calculate the correlation coefficient between the two images. We use positive constants *C*_1_, *C*_2_ and *C*_3_ to avoid null denominators [26]. Finally, we calculated distorted residuals of the incident pattern in the vertical orientation (**Fig. 5C**), which was equal to the intensity of the total distorted light field at the focal plane minus the intensity of the incident pattern. Thus, the spatial localization of the distorted pattern is consistent with the scattered light field.

**Fig. 5.**
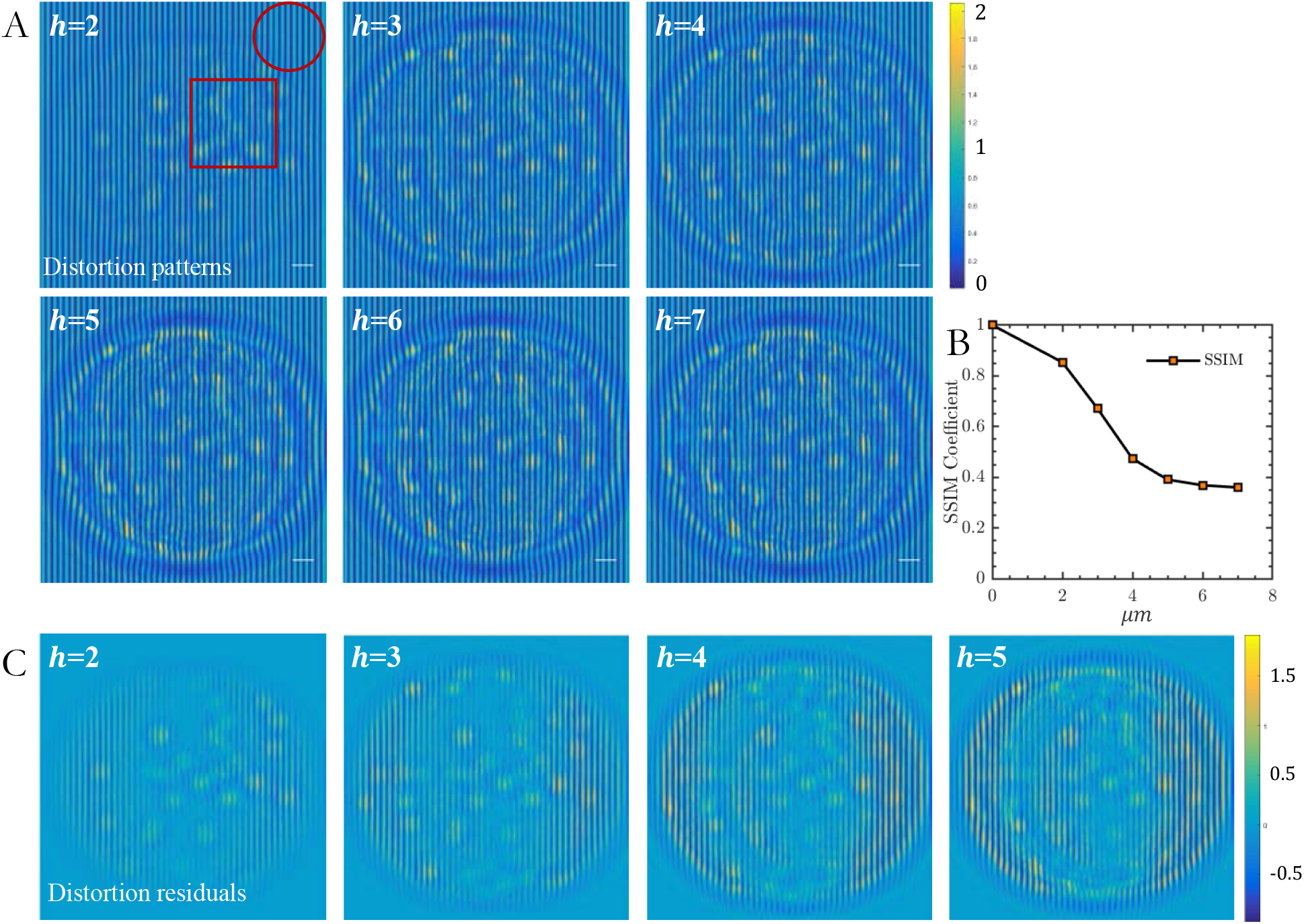
Distortions of the illumination patterns in the focal plane of imaging. (A) Intensities of illumination pattern with scattered distortions of a single phase in the vertical orientation in different sample thicknesses. (B) SSIM of the distorted patterns with *h* of 2 - 7 *μm* compared to the distortion-free pattern with *h* = 0. (C) Distortion residuals of the incident pattern in the vertical orientation, which are equal to the total optical field’s intensities at the focal plane minus the incident patterns. Scale bar: 2 *μm*.

SIM microscopy achieves improved resolution compared to a wide-field image, as observed from the comparison between the wide-field image and the SIM reconstruction image at *h* = 0 (**Fig. 6A**). However, because SIM microscopy illuminates the sample with structured patterns, reconstruction after image acquisition was required. If the illumination pattern is distorted during image acquisition, distortions will eventually form severe artifacts in the final SR image which is reconstructed by the commonly used Wiener-SIM [8] (**Fig. 6B**). At *h* = 0 *μm*, there are no reconstruction artifacts. However, striped or honeycomb artifacts begin to emerge in the reconstruction of objects of low spatial frequency when the illumination scattering occurs. For instance, at *h* = 2 *μm*, the artifacts are relatively weak, and the structure of the quadrilateral above, the circular area at the lower left and the rings at the lower right of the reconstructed image are relatively uniform and complete. As *h* increases, the contrast of the honeycomb artifacts becomes stronger, accompanied by a decrease in SSIM (**Fig. 6C)**. Even though some areas of illumination patterns are badly distorted (box in **Fig. 5A**), while other areas are less affected (circle in **Fig. 5A**), striped artifacts exist throughout the whole reconstruction image **(Fig. 6B**).

**Fig. 6.**
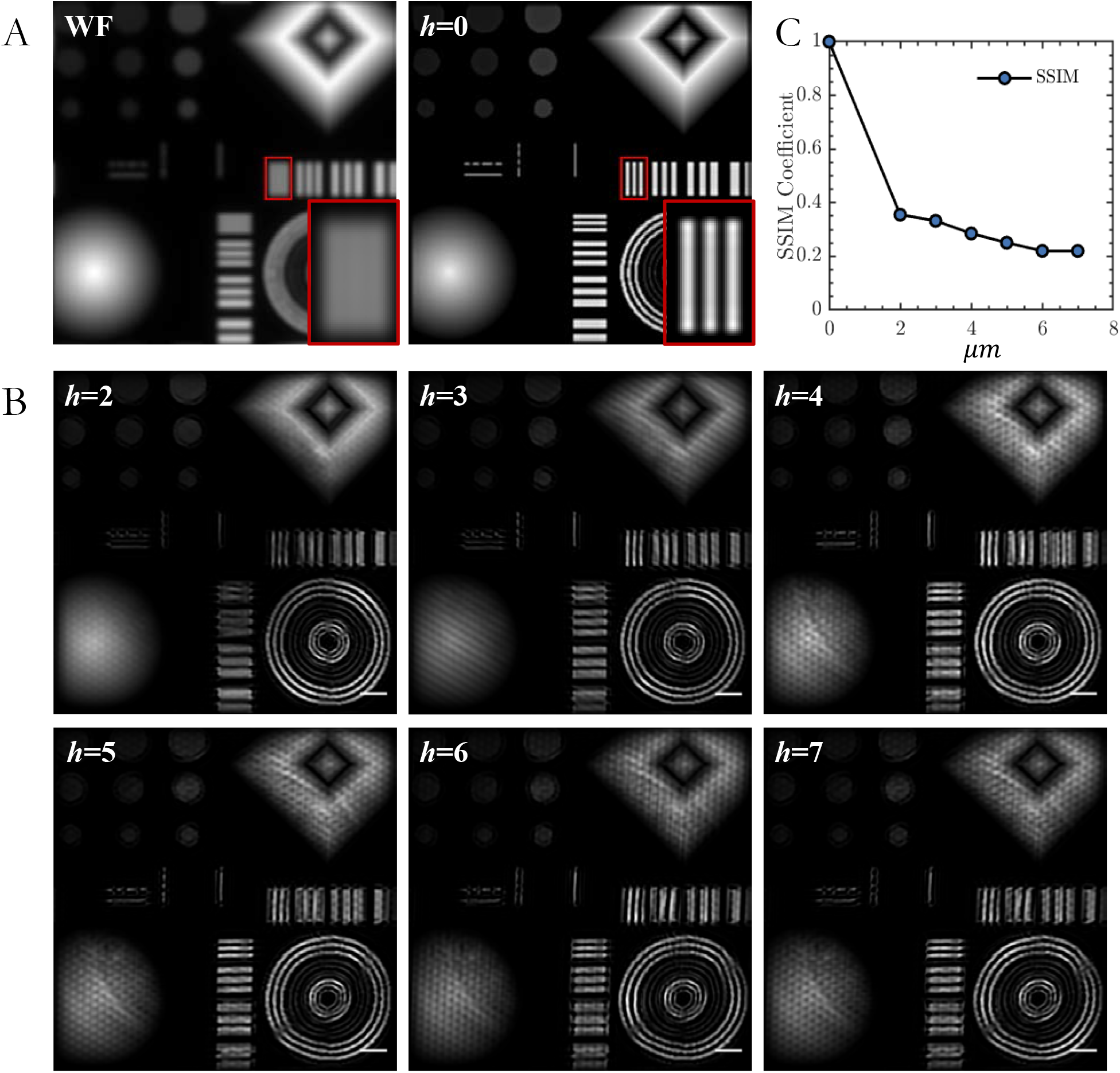
SIM reconstruction images with different *h*. (A) Wide-field image and SIM reconstruction image with *h* = 0. (B) Reconstruction results with different thickness distortion using commonly used Wiener-SIM. (C) SSIM index of the SIM reconstruction images with *h* of 2-7 *μm* compared to that with *h* = 0 *μm*. Scale bar: 2 *μm*.

The whole field of view was used to directly calculate global parameters (direct parameters) in Wiener-SIM, which might not match the heterogeneous distortion field and cause reconstruction artifacts. To distinguish artifacts caused by the scattering from those caused by the reconstruction, we have compared artifacts under the parameters determined under scatter-free conditions (*h* = 0) and parameters determined under scattered conditions (*h* > 0). These parameters exhibited significant variations as estimated under different *h*. For example, the deviation of the periodic *p*_*θ*_ in each orientation did not increase uniformly with the increase of thickness in scattered samples’ (**Table S1** and **Fig. S1**). With these estimated parameters, the reconstruction results and their effective OTFs (coupled modulation depth) are shown in **Fig. 6** and **Fig. S2**.

As compared to the SR images of direct parameters, reconstructing scattered objects (*h* > 0) using parameters determined from objects without scattering (*h* = 0, corrected parameters) always yielded better reconstructions (**Fig. 7** and **Fig. S3**), and demonstrated higher SSIM than the former. These data rather highlighted inaccurate direct parameters to be one major source of artifacts. However, even reconstructions using corrected parameters failed as imaging depth increased, which appeared to a contrary correlation for plane thickness and the SSIM of the reconstruction results (**Fig. S3**). In this sense, the scattering of illumination patterns also significantly influences the reconstructed SR images.

**Fig. 7.**
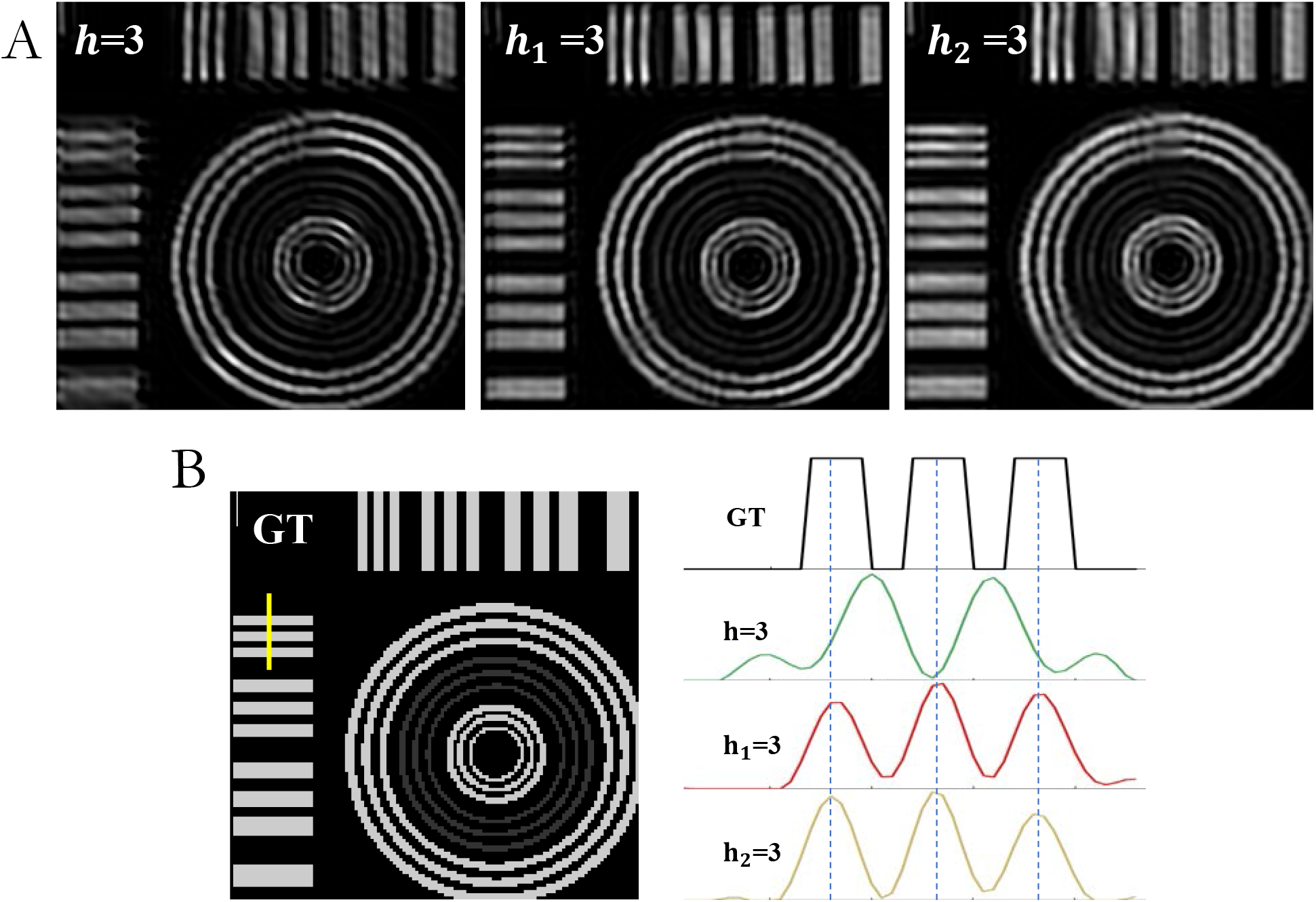
Result of direct reconstruction, subregion reconstruction and corrected parameters reconstruction when *h* is 3 *μm*. (A) The SR images reconstructed by three different parameters, where *h*, *h*_1_, *h*_2_ represents direct, subregion and corrected parameters reconstruction, respectively. (B) Comparison of these three reconstructions with the ground-truth image (GT) along the yellow line.

Locally estimating parameters of illumination pattern in a sub-region destroyed by the scattering may yield better reconstruction than corrected parameters. To test this hypothesis, we used parameters locally determined from a relatively uniform sub-region (*h* = 3) to reconstruct this region. The reconstructed sub-region was superior to that extracted from the whole image which is obtained with direct parameters, and is slightly better than that obtained with corrected parameters (**Fig. 7**). However, because the whole image field is scattered to different extents laterally and axially, the same uniform sub-region became non-homogeneous at a different image plane. Thus, parameters locally determined from other planes (*h* ≥ 5) deviated significantly from corrected parameters, and concentric rings in the reconstructed image were largely distorted (**Fig. S4**). These data pinpoint the challenge: how to non-biasedly sort out relative uniform subregions in three dimensions, and accurately estimate local parameters with smaller areas and fewer pixels available.

## Conclusions and discussion

In the simulation experiments, when the illumination light of SIM is scattered as it travels through the sample with an inhomogeneous refractive index, uniform fluorescent objects become corrupted with tweaked or honeycomb artifacts. These data are consistent with the live-cell SR experimental data, where shaped artifacts are often obvious in the observed organelles and structures deep inside the cell using SIM microscopy [7,27–29]. These data suggest that our simulation results may be used to guide artifact suppression in real live-cell SR experiments. Because of distortion, the illumination patterns are spatially variant over the sample. While it seems improper to directly reconstruct the whole field of view with direct parameters, we showed here that corrected parameters yielded better results and fewer artifacts than direct parameters (Fig. S3). Therefore, by recording the scattered images as well as distributions of RIs in 3D, it may be possible to reversely derive the original signals from scattering, and determine the corrected parameters for reconstruction with fewer artifacts. Alternatively, for the given RI distributions within the thick sample corresponding to the SIM raw data, we can obtain the specific distribution of the actual illumination pattern distortion caused by the thick sample’s scattering. Similar to MAP-SIM [30], we may be able to build an optimization model that substitutes the cost function with the actual distorted patterns. By using the iterative optimization method to obtain reconstructed images of better quality, we may reduce artifacts caused by the illumination light scattering in the future.

Here we only simulate the scattering of incidence. The scattering of the fluorescence emitted by the sample on the detection path will also introduce aberrations and distortions to the detected PSF[17]. With the 3D refractive index distribution measured by the ODT, it may be possible to correct the distortions due to the aberrant illumination pattern during fluorescence excitation and handle the harmful effects of the scattering of emitted fluorescence. Our goal is to reduce artifacts and achieve ideal, high-fidelity SR-SIM imaging in live cells.

## Additional Information

### Data Accessibility

All datasets generated for this study are included in the article material.

### Authors’ Contributions

Y.M., F.F., and H.M carried out the experiments. Y.M. performed the data analysis and drafted the manuscript. J.F. and L.C. conceived and designed the study, and wrote the manuscript. All authors read and approved the manuscript.

### Disclaimer

The authors confirm that there are no conflicts of interest.

