## Supplementary for "Structured illumination microscopy artifacts caused by illumination scattering"

Table.S1 | Estimated illumination parameters

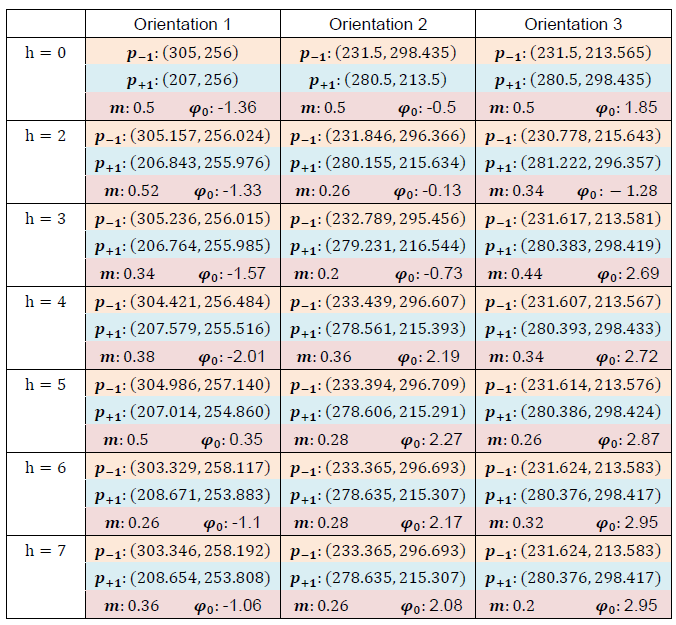

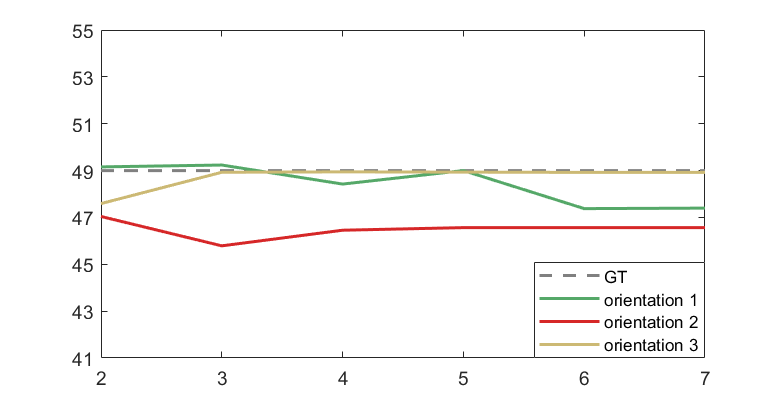

Fig. S1 | Length of illumination periodic $\boldsymbol{p}_{\theta}$ in each orientation of direct parameters.

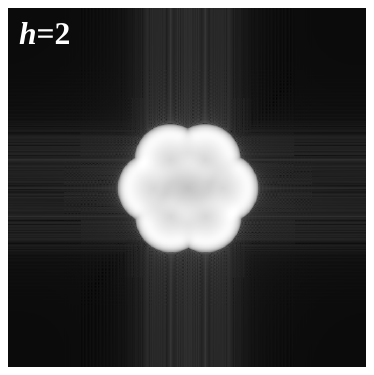

Fig. S2 | The effective OTF of direct parameters (coupled modulation depth).

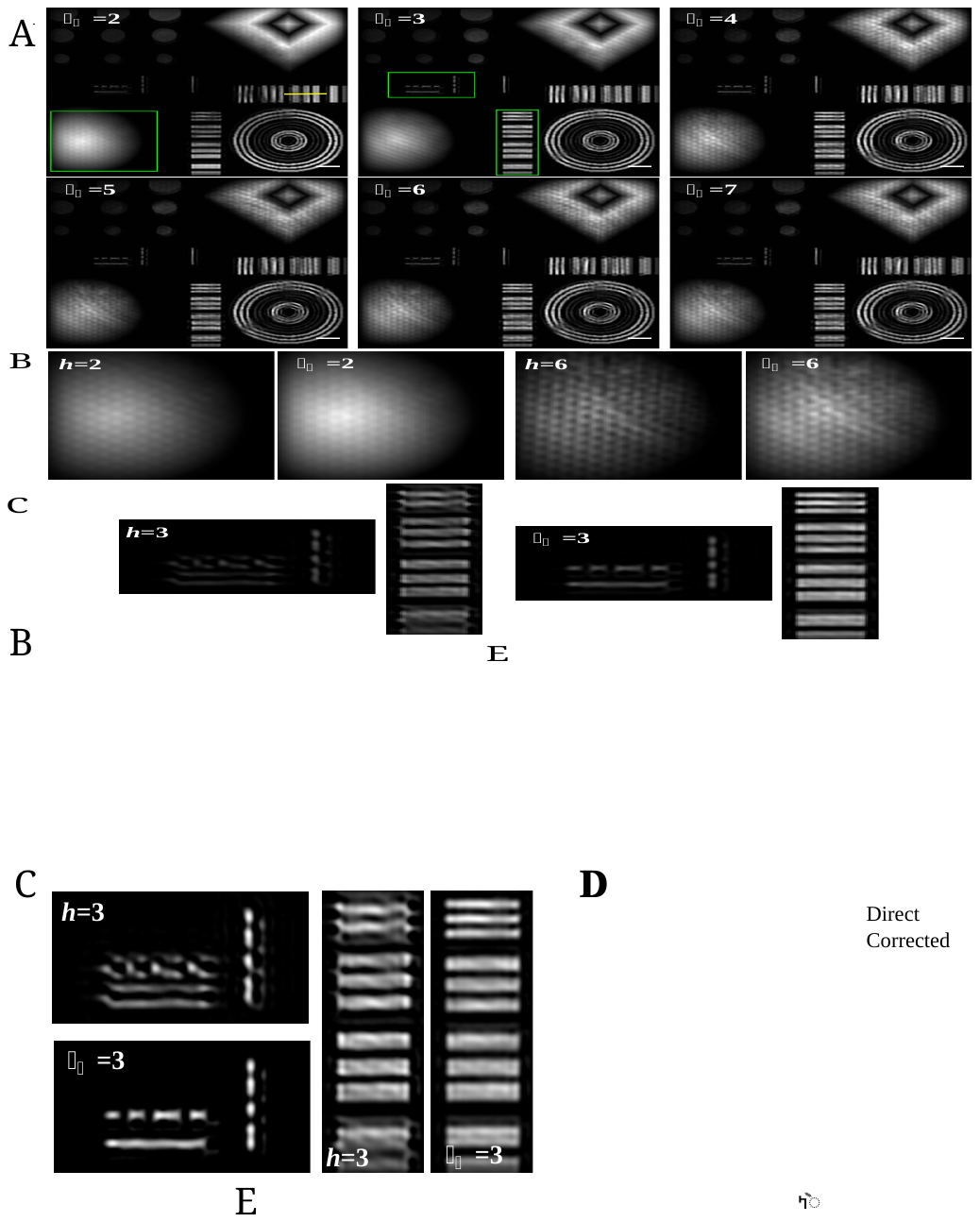

Fig. S3 | Results of parameter-corrected reconstruction. (A) The parameter-corrected reconstruction results (*h* > 0), which used parameters determined from objects without scattering (h = 0). $h$ and $h_{2}$ represents the results of direct parameters and corrected parameters respectively. (B) Comparison between the results with direct parameters and those with corrected parameters at the green square ROI in (A). (C) Comparison between the results with the direct parameters and those with corrected parameters at the green rectangle ROI in (A). The area that was mistakenly reconstructed into two segments can be restored to one line segment with corrected parameters. And the uniformity of the transverse wide line segments become better. (D) SSIM of the SR images reconstructed with the direct parameters and corrected parameters at different thicknesses. The solid blue line is the SSIM of the direct parameters result, and the dotted red line is the SSIM of the corrected parameters. (E) The intensity of the parameter-corrected reconstruction at different *h* along the yellow line in (A). Scale bar: 2 $\mu m$*.*

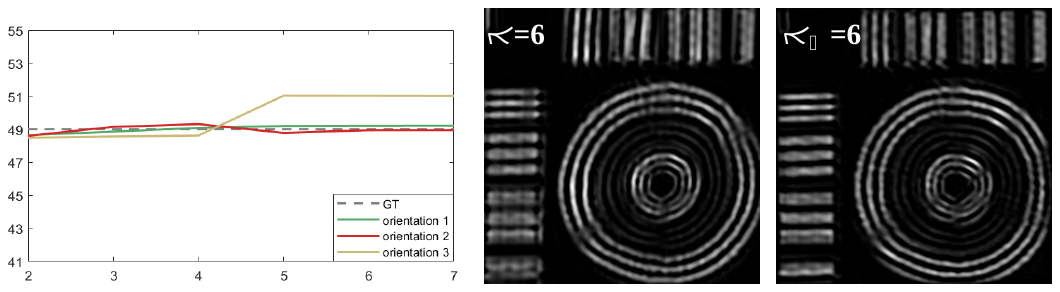

Fig. S4 | Length of $\boldsymbol{p}_{\theta}$ in each orientation of subregion reconstruction and the results when *h* is 6 $\mu m$. $h$ and $h_{1}$ represents the result of direct parameters and subregion reconstruction respectively.
